# HSDatabase – a database of highly similar duplicate genes from plants, animals, and algae

**DOI:** 10.1101/2022.08.01.502183

**Authors:** Xi Zhang, Yining Hu, David Roy Smith

## Abstract

Gene duplication is an important evolutionary mechanism capable of providing new genetic material, which can help organisms adapt to various environmental conditions. Recent studies, for example, have indicated that highly similar duplicated genes (HSDs) are involved in adaptation to extreme conditions via gene dosage. However, HSDs in most genomes remain uncharacterized. Here, we collected and curated HSDs in nuclear genomes from a diversity of species and indexed them in an online, open-access sequence repository called HSDatabase. Currently, this database contains 117,864 curated HSDs from 40 eukaryotic genomes, and it includes information on the total HSD number, gene copy number/length, and alignments of gene copies. HSDatabase also allows users to download sequences of gene copies, access genome browsers, and link out to other databases, such as Pfam and KEGG. What’s more, a built-in Basic Local Alignment Search Tool (BLAST) option is available to conveniently explore potential homologous sequences of interest within and across species. HSDatabase is presented with a user-friendly interface and provides easy access to the source data. It can be used on its own for comparative analyses of gene duplicates or in conjunction with HSDFinder, a newly developed bioinformatics tool for identifying, annotating, categorizing, and visualizing HSDs.

**Database URL:** http://hsdfinder.com/database/

## Introduction

Gene duplication is a near-ubiquitous phenomenon throughout the eukaryotic tree of life (1), one that can be advantageous or disadvantageous, depending on the circumstances. For example, under certain conditions, it can be detrimental for an organism to retain highly similar expressed genes (2). Thus, with notable exceptions, it is relatively rare for species to maintain duplicate genes encoding the same functions (3). Nevertheless, it is becoming more apparent that in some situations the generation and maintenance of highly similar duplicate genes (HSDs) is possible, particularly for genes encoding products that are in high demand, such as histones or ribosomal proteins (4). Indeed, there are many examples suggesting that stress response genes, sensory genes, transport genes, and genes that have a metabolism-related function are likely to be fixed as duplicated copies given specific environmental conditions (5).

Recently, Zhang *et al*. (6) revealed that hundreds of HSDs, involved in diverse cellular processes, are maintained in the psychrophilic Antarctic green alga *Chlamydomonas* sp. UWO241, which was recently renamed *Chlamydomonas priscuii* (7). It is believed that these HSDs are aiding its survival via gene dosage (8). Unfortunately, the HSDs from most other eukaryotic genomes, particularly those of algae, remain uncharacterized. This is partly because the experimental methods for identifying HSDs are time-consuming and labor-intensive. Many of the available bioinformatics tools for characterizing gene duplicates are limited by their designs (e.g., they only identify paralogs) or their specificity (e.g., only identify retrocopies or co-localised duplicates) (9-13). Consequently, we recently developed a web-based tool called HSDFinder that can identify HSDs in eukaryotic genomes with high accuracy and reliability (14). For example, HSDFinder predicted 336 and 265 HSDs in the psychrophilic green algae UWO241 and *Chlamydomonas* sp. ICE-L (15), respectively, which is consistent with other experimental data (8). By applying HSDFinder to a variety of other species (16), we predicted and cataloged thousands of HSD candidates, which are now curated and documented in a new online repository called HSDatabase. Currently, it houses 117,864 HSDs from 40 eukaryotic species, with a focus on green algae, animals and land plants.

Here, we briefly introduce the general features as well as the procedures and principles for collecting data from HSDatabase. In short, HSDatabase contains information on HSD number, gene copy number, and gene copy length. Additionally, the protein functional domains and associated pathways of the HSDs can be retrieved from the Kyoto Encyclopedia of Genes and Genomes (KEGG) and InterProScan. A built-in BLAST-search option is also provided, allowing users to conveniently explore potential homologous sequences of interest within and across species. HSDatabase also provides data on a range of parameters and questions about gene duplicates, such as number of HSD per Mb, the most commonly conserved domains among HSDs, and the functional categories of HSDs. It is our hope to build a comparative analysis framework across species, especially for best-assembled eukaryotic genomes from species living in extreme environments, to better understand the role of gene duplication in adaptive evolution.

## Materials and Methods

### Database collection

HSDs were identified in 40 well-assembled nuclear genomes from diverse model species, including from plants and algae (e.g., *Arabidopsis thaliana, Chlamydomonas reinhardtii, Fragilariopsis cylindrus*, and *Zea mays*) and animals (e.g., *Drosophila melanogaster, Homo sapiens*, and *Mus musculus*). We selected mostly animal and plant genomes because of their high-quality assemblies. The genome assemblies of the selected species are retrievable from the National Center for Biotechnology Information (NCBI) (17) (Table 1). As displayed in a taxonomic tree (Figure 1), the selected species are highlighted with different kingdom colors based on NCBI Taxonomy labels (18, 19). The HSDs, which are represented by gene copies with nearly identical lengths and similar gene structures, were identified using HSDFinder (14). The identification method is based on all-against-all BLASTP analyses (20), which perform protein sequence similarity searches with specific uniform homology assessment metrics, e.g., E-value cut-off ≤1e-10, amino acid pairwise identity ≥90%, and amino acid aligned length variance ≤10 (note, the short form for these metrics is “90%_10aa”). Additionally, the putative HSDs are expected to have similar structural information, such as matching protein family (Pfam) domains (21), corresponding InterPro annotations (22), and/or nearly identical conserved residues. The InterProScan tool (23), which is an integrated platform for protein signatures, was used to collect the structural information on the HSDs. With the all-against-all BLAST and InterProScan results (tab-delimited files) available, they can be fed easily into the HSDFinder tool to generate HSD candidates in an 8-column tab-delimited file (Figure 2A). HSDs candidates can be identified via parsing the BLAST all-against-all protein similarity search result via the homology metrics: amino acid pairwise identity and amino acid aligned length variance. To collect and curate the data in HSDatabase, we performed a series of combo thresholds for filtering putatively functional gene copies.

**Table 1.**
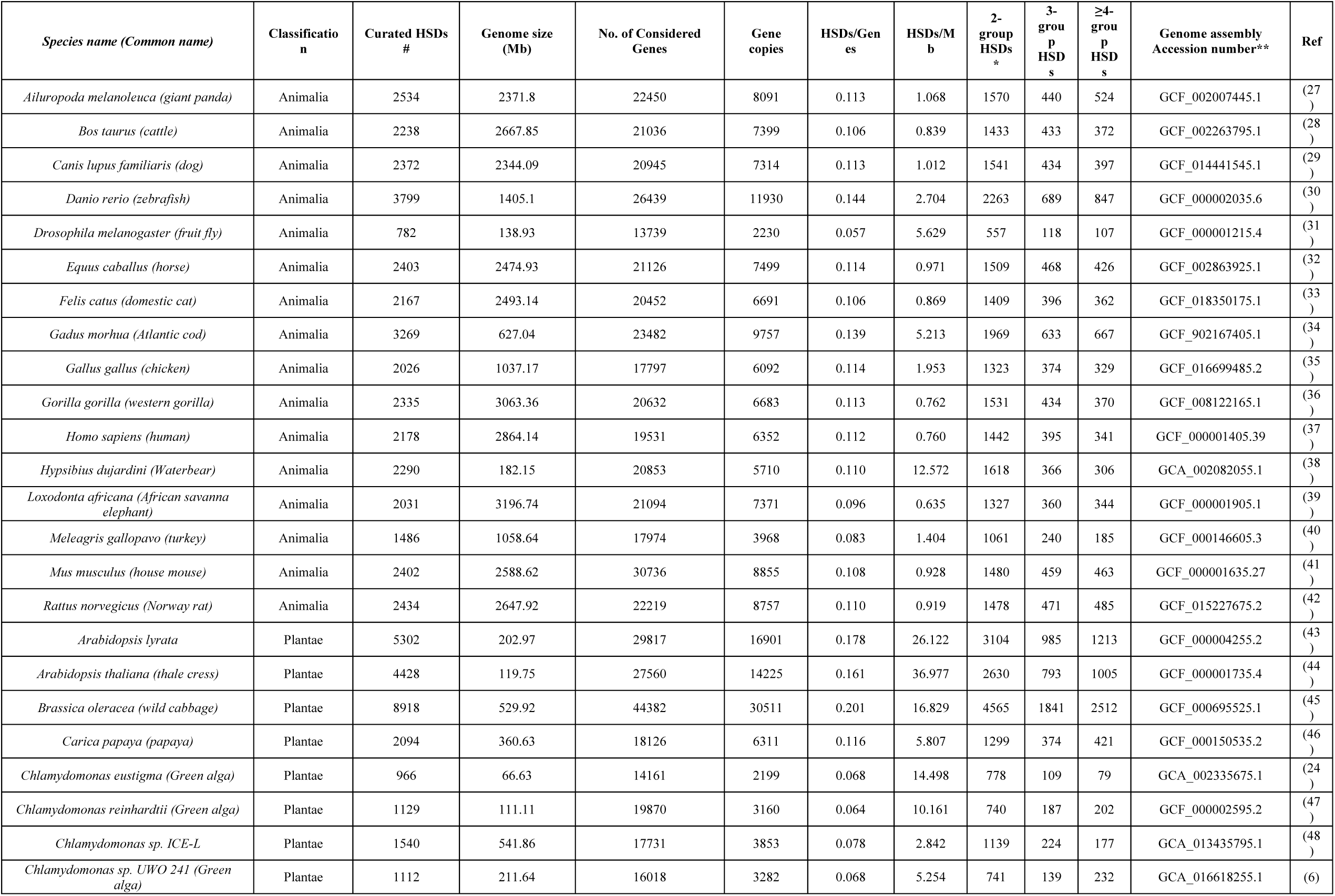
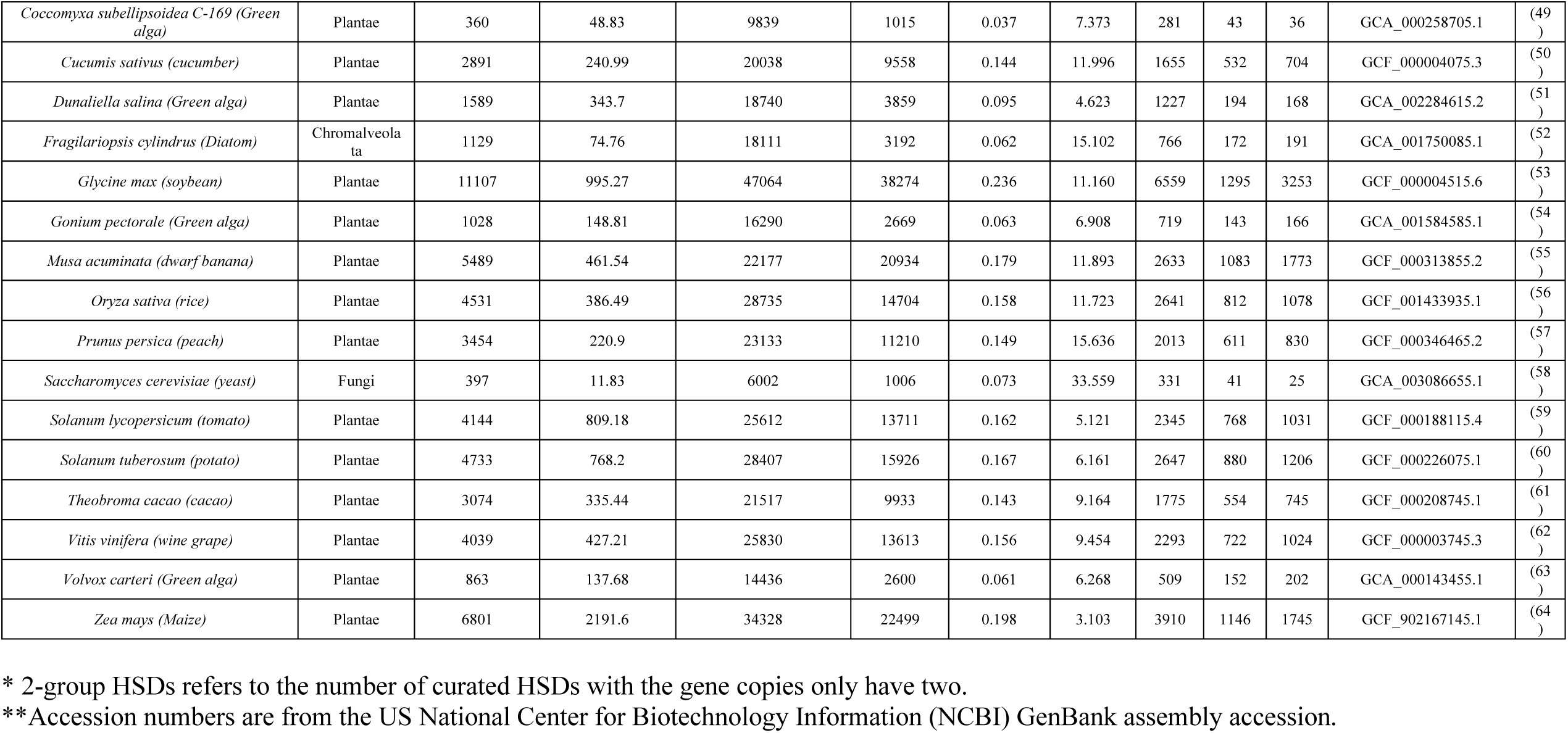
Summary statistics of the curated HSDs in the selected genomes from HSDatabase.

**Figure 1.**
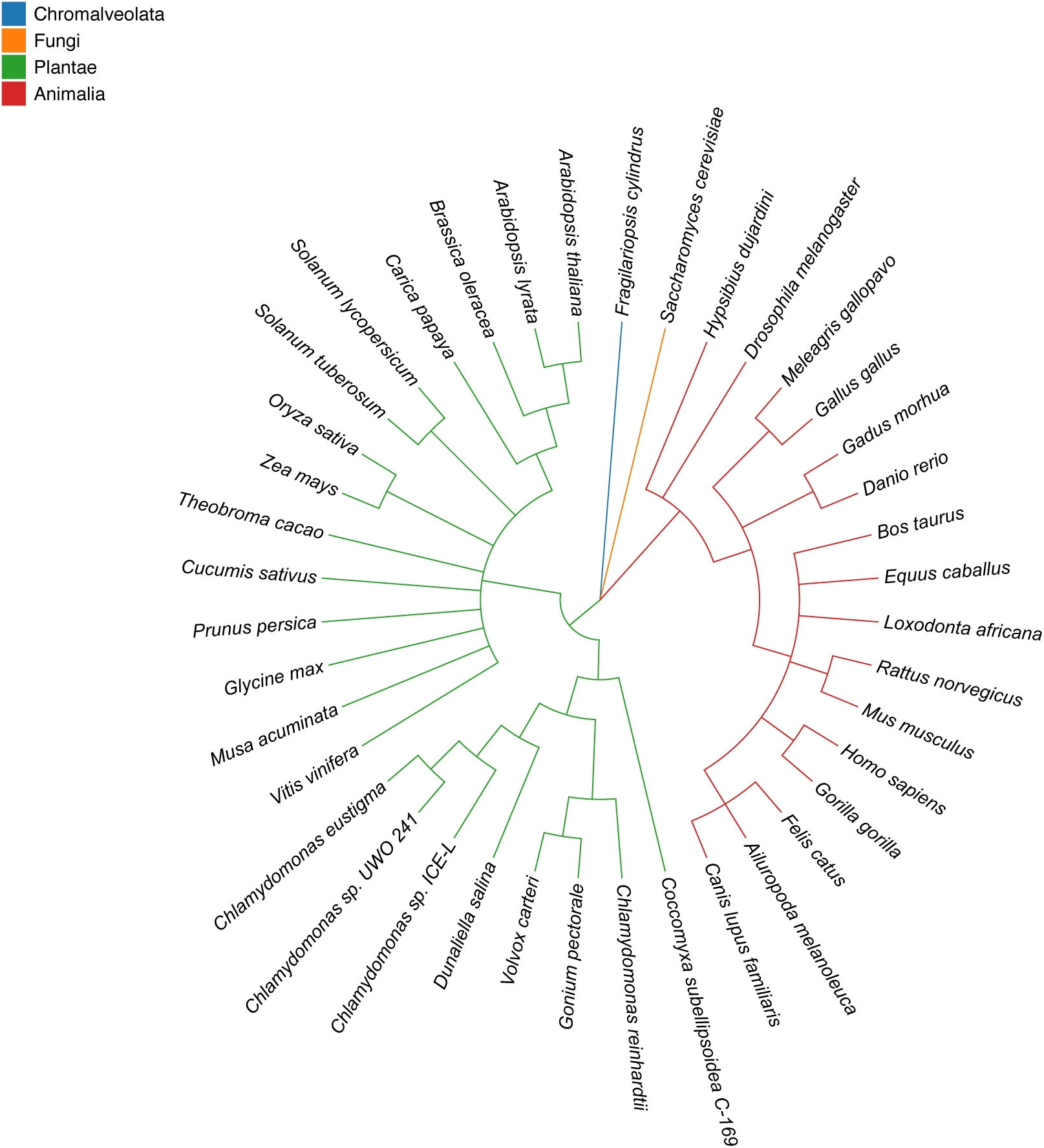
Taxonomic tree of 40 eukaryotic species in four highlighted categories. Chromalveolata, Fungi, Plantae and Animalia are in blue, yellow, green and red, respectively. The tree topologies were inferred by Taxonomy Common Tree from NCBI (https://www.ncbi.nlm.nih.gov/Taxonomy/CommonTree/wwwcmt.cgi).

**Figure 2.**
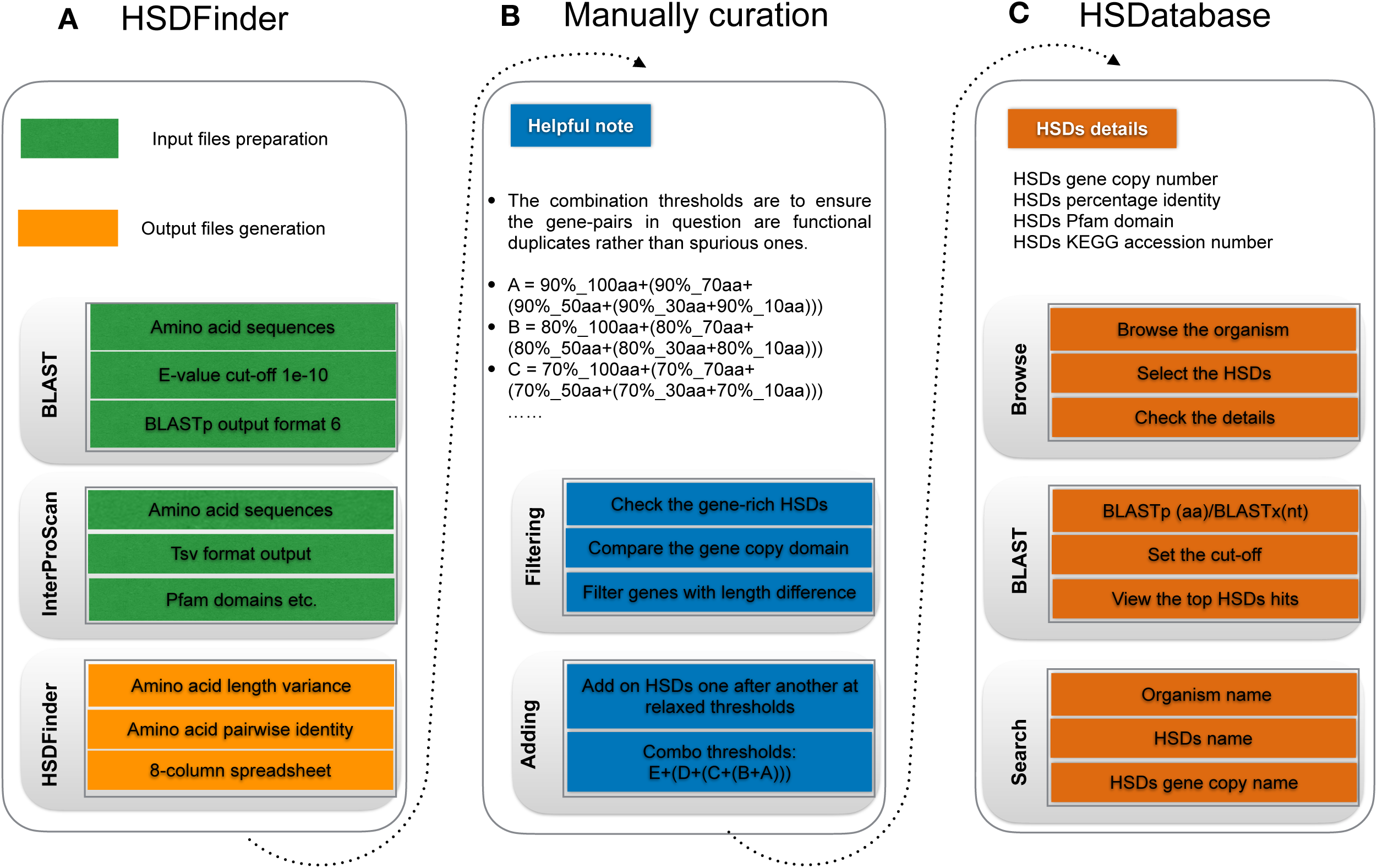
The workflow of HSDatabase. (A) Steps for using HSDFinder to collect candidate HSDs. (B) Manual curation of HSDs via filtering and adding new HSD candidates before putting into HSDatabase (C) The steps of accessing the HSDs records in HSDatabase, including browsing via the organism names, BLAST query sequences against database, and searching through the HSD and gene copy IDs.

### Database curation

Prior to uploading data into HSDatabase, we curated HSD candidates by filtering for redundancy and adding the newly curated HSDs (Figure 2B). For genes that have alternative protein products, we maintained the longest gene isoform to reduce redundancy. Since highly similar gene copies are grouped together as HSDs based on a simple transitive link between the remaining genes (14), it is possible for some genes enriched with duplicates to form a mega HSD group with varied gene copy lengths, especially for those encoding histones, ribosomal proteins, and retro-transcriptases. Moreover, some gene copies might appear multiple times causing redundancy among different HSD groups, which is because the BLAST algorithm limits the maximum target hits by default. In these cases, we manually curated the HSD groups, minimizing redundant gene copies.

Since the similarity of duplicated genes within and among genomes can vary significantly, we added newly curated HSDs to the database using a combination of thresholds to acquire a larger dataset of HSD candidates. We added the HSDs candidates one after another at different homology assessment metrics (i.e., HSDs identified at more relaxed thresholds were treated more strictly than those found using more conservative thresholds) (Figure 2B). For example, HSDs identified at a threshold of 90%_30aa were added on to those identified at a threshold of 90%_10aa (denoted as “ 90%_30aa+90%_10aa”); any redundant HSDs candidates picked out at this combo threshold were removed if the more relaxed threshold (i.e., 90%_30aa) had the identical genes or contained the same gene copies from the stricter cut-off (i.e., 90%_10aa). Moreover, any HSDs candidates pinpointed at the combo threshold (90%_30aa+90%_10aa) were removed if the minimum gene copy length was less than half of the maximum gene copy length for each HSD, or if HSD candidates had gene copies with incomplete conserved domains (i.e., different number of Pfam domains). After filtering the combo threshold at (90%_30aa+90%_10aa), we added on a more relaxed threshold 90%_50aa (i.e., 90%_50aa+(90%_30aa+90%_10aa)) and then carried out the same HSD candidate removal/filtering process. To minimize the redundancy and to acquire a larger dataset of HSD candidates, we processed each selected species with the following combination of thresholds: E + (D + (C + (B +A))).

A = 90%_100aa+(90%_70aa+(90%_50aa+(90%_30aa+90%_10aa)))
B = 80%_100aa+(80%_70aa+(80%_50aa+(80%_30aa+80%_10aa)))
C = 70%_100aa+(70%_70aa+(70%_50aa+(70%_30aa+70%_10aa)))
D = 60%_100aa+(60%_70aa+(60%_50aa+(60%_30aa+60%_10aa)))
E = 50%_100aa+(50%_70aa+(50%_50aa+(50%_30aa+50%_10aa)))

### Database implementation

The database was built with the Django 3.0.5 web framework (https://www.djangoproject.com/), and all data were stored in an SQLite 3.36.0 database (https://www.sqlite.org/index.html) on an Amazon web server. Webpage templates used Bootstrap framework (https://getbootstrap.com/), D3.js (https://d3js.org), jQuery (http://jquery.com), and Bootstrap Table (https://bootstrap-table.com/) libraries to establish a user-friendly, front-end interface. On the browse page, NCBI’s Sequence Viewer 3.44.0 (https://www.ncbi.nlm.nih.gov/projects/sviewer/) was employed to build a fast and scalable genome browser.

## Results and Discussion

### Database content and analysis

HSDatabase is built using a relational database (MySQL) allowing rapid retrieval of data and making resources easily maintainable. One entry corresponds to one eukaryote genome. The genomes can be accessed via the organism table or the taxonomic tree. The genome entry is then split into various subcategories of HSD entries. The database access is via a web interface based on python script and provides various ways to search for HSD entries, including species name, unique HSD IDs, and gene copy IDs.

Using HSDFinder (16), we collected and curated 117,864 HSDs and 379,844 gene copies from 40 well-assembled nuclear genomes from diverse model species (Table 1). Various green algae were included because of our specific interest in algal genomics and also because of their relatively small genome sizes and gene numbers. For example, the acidophilic green alga *Chlamydomonas eustigma* is known to have large numbers of gene duplicates in its nuclear genome, including 10 copies of genes encoding arsenate reductase (ArsC) and 20 copies of genes encoding glutaredoxin (Grx) (24). Similarly, the psychrophilic green alga *Chlamydomonas* sp. ICE-L contains multiple copies of genes encoding carotene biosynthesis-related protein (CBR) and Lhc-like protein (Lhl4) (25). These data are consistent with our identifying large numbers of HSDs in *C. eustigma* (276) and ICE-L (265) (Table 1), suggesting a potential adaptative role of gene duplication under different extreme environmental conditions. Compared to animals, the investigated land plants had higher detected numbers of HSDs, as well as higher levels of HSDs/Mb and HSDs/genes values (Table 1). For example, the HSDs/Mb values for *A. thaliana, Arabidopsis lyrate*, and *D. melanogaster* are 37.0, 26.1, and 5.6, respectively, whereas the average HSDs/Mb value among selected green algae is 8.2. The large difference in HSD number between land plants and green algae is partly a reflection of the explored land plants having diploid genomes, which can yield more gene duplicates via whole-genome duplication events compared to the effectively haploid green algal genomes. The number of 2-group HSDs (i.e., HSDs only contain two gene copies) represent the majority (>50%) of the total HSDs for each selected species.

As for the function of the detected HSDs, three green algal species with relatively large values of HSDs/genes were compared previously. Specifically, these algae can survive under different extreme environmental conditions, such as the Antarctic psychrophilic green algae UWO241 (0.068) and ICE-L (0.078) and the acidophilic *C. eustigma* (0.068) (Table 1). The identified duplicates are involved in a diversity of cellular pathways, including gene expression, cell growth, membrane transport, and energy metabolism, but also include ribosomal proteins (6, 14). Although HSDs for protein translation, DNA packaging, and photosynthesis are particularly prevalent, around 30% of the HSDs are hypothetical proteins without any Pfam domains.

### Database composition and usage

Information about specific HSDs and their associated gene copies for a given species can be obtained using one of two main approaches: i) the “Browse” and “Search” tabs, which are located on the menu bar at the top of the page; or ii) using protein or nucleotide sequences as queries to search against the database via BLAST (i.e., BLASTP or BLASTX). To categorize duplicated genes into a functional category, KEGG pathway schematics are available for each species.

#### Browse

By selecting the ‘Browse’ option from the main menu, users are offered three ways to explore their species of interest. First, they can simply click the organism name on the taxonomic tree containing the 40 species within HSDatabase. Secondly, users can select the tab for “Plantae, Fungi, and Chromalveolata”. Twenty-three species were categorized into a summary table documenting organism name, number of HSDs, species background information, GenBank accession links to the genome assembly, and reference links to PubMed. Thirdly, users can select the “Animalia” tab to see these same statistics for the seventeen animal species (Figure 3A).

**Figure 3.**
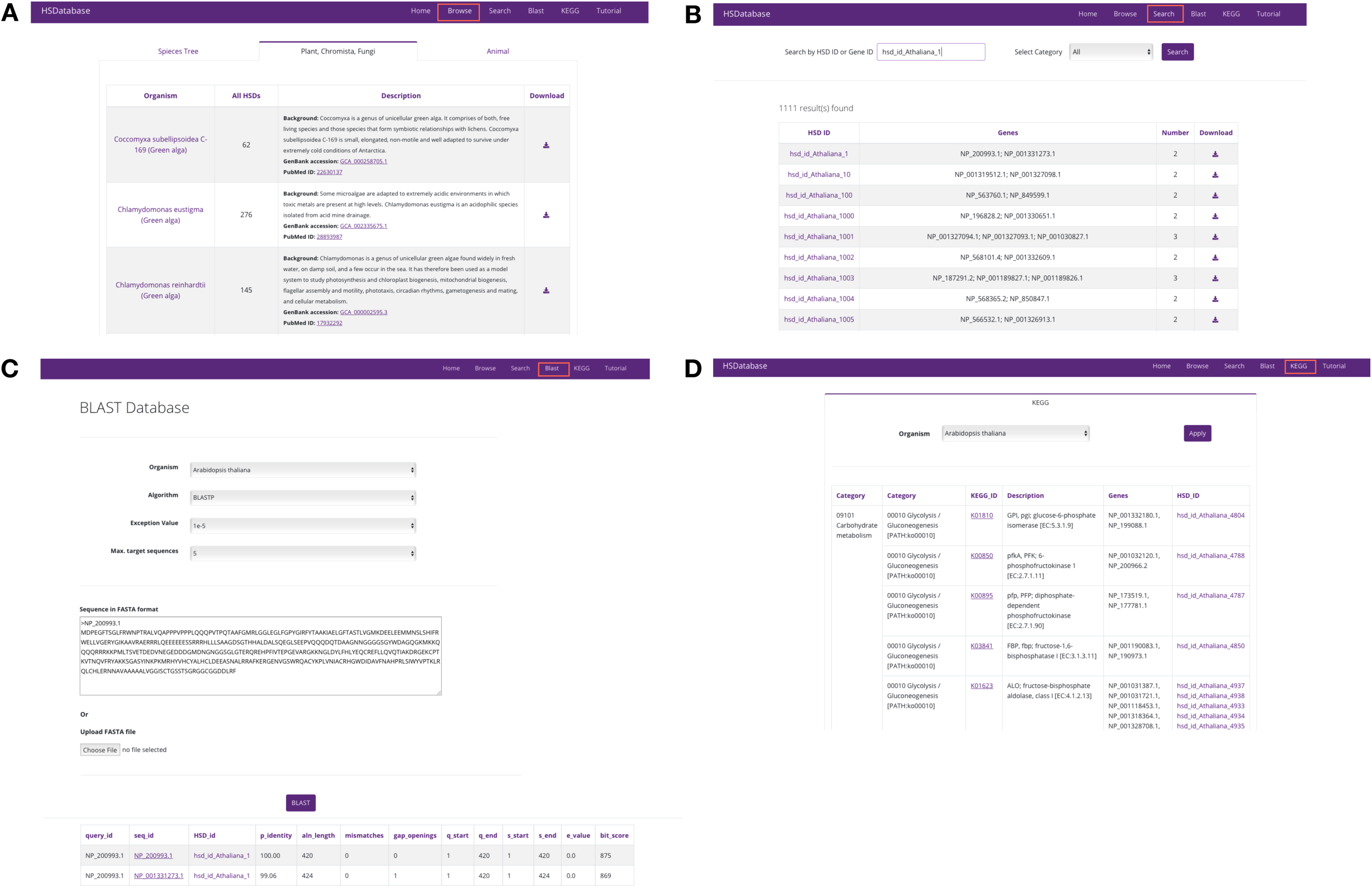
The screenshots of the HSDatabase interface. There are four main functions in the menu page: (A) Browse the database via the species entries; (B) Search the database via the HSDatabase unique ID (e.g., hsd_id_Athaliana_1) or gene ID (e.g., NP_200993.1); (C) Use BLAST to search the database via the protein sequence in FASTA format; and (D) Categorize the gene copies and HSDs under the KEGG pathway functional categories.

The selection of a specific species from the browse page leads to the respective HSDs summary page (Figure 4A). This page provides the total number of HSDs, unique HSD IDs, gene copy GenBank IDs, number of gene copies, and access to the data download function. Choosing one of the HSD ID entries, brings up the basic information and features for a detected HSD. For example, the page provides details on the various gene copies for a unique HSD, including the GenBank link, the sequence length, the Pfam domain ID, Pfam domain description, and the InterPro database ID and description (Figure 4B). Clicking on the genome browser tab brings up the visualization of a specific gene copy in a built-in NCBI genome browser (Figure 4C). The FASTA sequence tab provides the option to download the sequence data (Figure 4D). The “alignments and identity%” tab provides the gene copy alignments and percentage identity matrix created by the built-in tool Clustal v2.1 tool (26) (Figure 4E).

**Figure 4.**
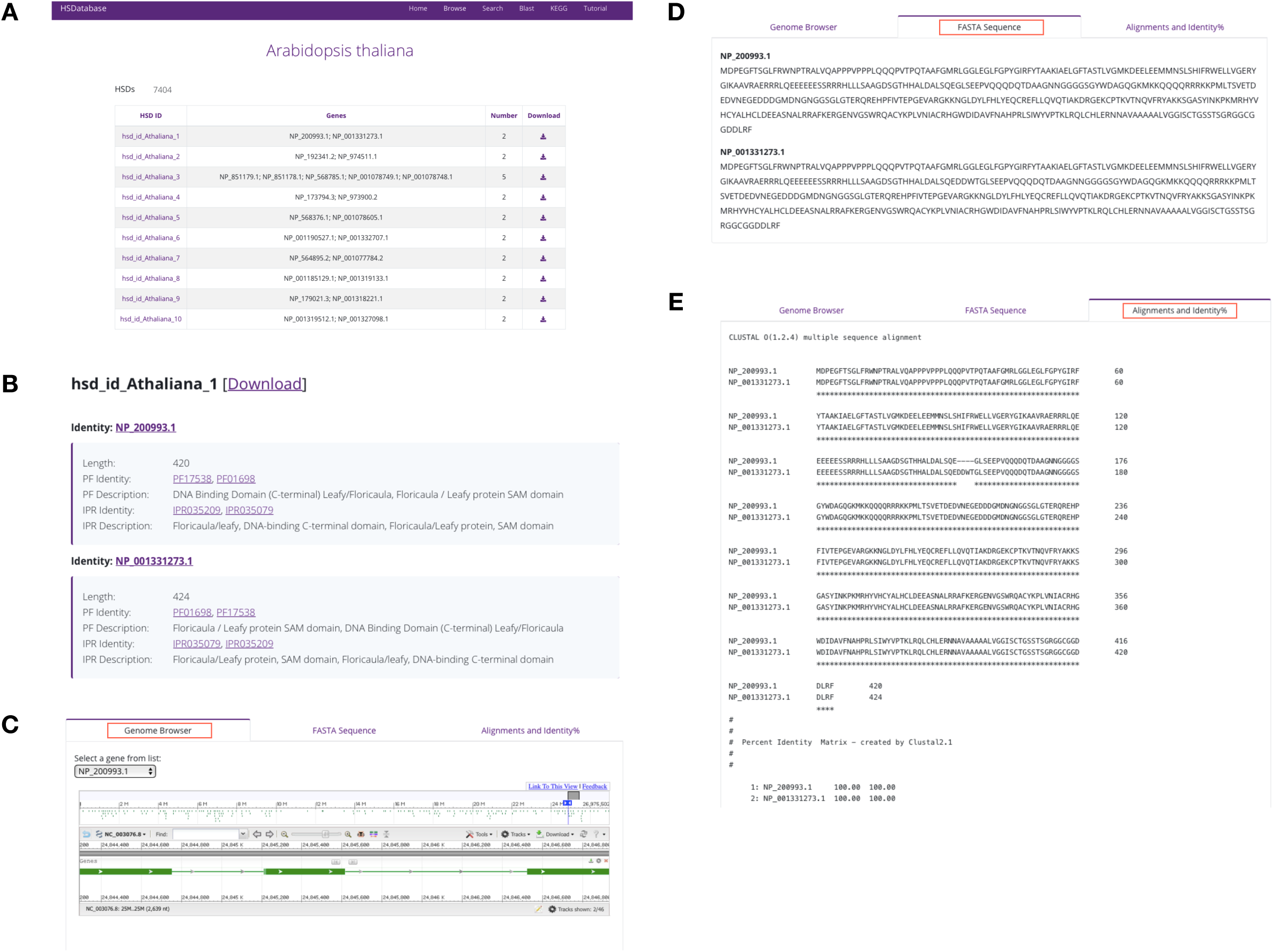
The summary of database information in a selected species. (A) HSDs collected in a table of a selected species (B) The basic information of the unique HSD ID, gene copy ID, and the associated links to Pfam domains and InterPro databases. (C) The links of gene copies to genome browsers. (D)The FASTA sequence downloads of gene copies. (E) The alignments and percentage identities of the gene copies.

#### Search

By choosing the search option from the main menu, users can search unique HSD IDs or gene copy IDs from the database (Figure 3B). Users can also set the selection categories to limit the search results, which can improve the search efficiency. After activation of the search button, 30 results per page will be displayed (Figure 3B). Users can navigate through the results page or download specific HSD entries. The search results are summarized in a four-column table, including HSD name, gene copy name, number of gene copies, and the external download link to the output data (tab-delimited file). As described above in the Browse section, the data file includes various summary statistics on the HSDs (Figure 4B).

#### BLAST

The BLAST tool bar allows users to input a nucleotide or amino acid sequence (in FASTA format) to conduct a sequence similarity search using BLASTX or BLASTP. Users can specify the species against which the BLAST search will be performed. The e-value and maximum target sequence of results can also be adjusted, but all other parameters remain as defaults and cannot be changed (Figure 3C). The BLAST search output result is in standard 13-column tabular format, including the linkable query sequence ID and HSD ID, percentage identity, aligned length, and all other BLAST tabular output values. The most similar sequence targets are arranged at the top.

#### KEGG

The KEGG page of HSDatabase details the associated KEGG pathways of the HSD gene copies for 40 species. To browse one species, users can simply select the organism’s name. The 6-column table details the gene copies and HSDs under KEGG functional categories (Figure 3D). Gene copies involved in the same KEGG pathway are detailed with the first KEGG category (e.g., Carbohydrate metabolism), the secondary category (e.g., Glycolysis / Gluconeogenesis), and the KEGG pathway function description (e.g., ENO, eno; enolase). The KEGG ID (e.g., K01689) can also be linked to the external KEGG database with more detailed information.

### Future direction and limitation

Now that HSDatabase is publicly available, the next logical step is to analyze duplicate genes across a broader range of species, which we plan on doing in the coming months. Currently the database includes various straightforward statistics (e.g., number of HSD per Mb,), but there are many other statistics that can be added to the database, including information on differential expression levels among duplicates, for instance, as well as data on the rates of synonymous and nonsynonymous substitutions (dN/dS rates). The biggest challenge moving forward will be determining an appropriate threshold for accurately predicting HSDs. For example, how will we need to adjust the metrics (amino acids pairwise identity, amino acids aligned length variance, etc.) to find as many bona fide HSDs as possible? Presently, there is no standard golden cut-off for identifying HSDs and there might never be one as there a multitude of forces, including lineage/genomic specific ones, that can impact the accuracy of the identification metrics. This is why users can employ different parameters in the HSDFinder tool (from 30% to 100% amino acids pairwise identity or from within 0 aa to 100 amino acid aligned length variances). In our case, we use a series of combination thresholds to add on newly curated HSDs. In the future, our goal is to guide users to species-specific thresholds and deposit more diverse eukaryotic group species into HSDatabase.

### Conclusions

With the decreasing cost of next-generation sequencing, biologists are dealing with ever larger amounts of data. However, many bioinformatics software suites require considerable knowledge of computer scripting and microprogramming. To facilitate the understanding and analysis of gene duplication in nuclear genomes, we developed HSDatabase, which currently contains 117,864 HSDs from 40 well-assembled eukaryotic genomes. In conjunction, with HSDatabase, we designed the HSDFinder tool, which can efficiently identify duplicated genes from unannotated genome sequences by integrating the results from InterProScan and KEGG. HSDatabase aims to become a useful platform for the identification and comprehensive analysis of HSDs in eukaryotic genomes, which will aid research into the mechanisms driving genome adaptation. In the future, the database will be updated by taking into account more scientific discoveries in the field of gene duplication.

## Data Availability Statement

The datasets of eukaryotes supporting the conclusions of this article are available from JGI (https://phytozome.jgi.doe.gov/pz/portal.html) or NCBI (https://www.ncbi.nlm.nih.gov) database.

## Author Contributions

The study was conceptualized by XZ and DRS. The data were analyzed by XZ and YNH implemented the HSDatabase website. XZ and DRS drafted the manuscript and all authors commented to produce the manuscript for peer review.

## Acknowledgments

This work was supported by a Discovery Grants from the Natural Sciences and Engineering Research Council of Canada (NSERC) to DRS. We want to especially thank the editors and reviewers for their professional comments that greatly improved this manuscript.

## Supplementary Material

The predicted HSDs of 40 eukaryotes are documented in HSDatabase, which can be accessed via http://hsdfinder.com/database/. The HSDFinder source code has been deposited at https://github.com/zx0223winner/HSDFinder. The web server of HSDFinder is freely available at http://hsdfinder.com.

